# The SOCS1 KIR and SH2 domain are both required for suppression of cytokine signaling *in vivo*

**DOI:** 10.1101/2022.12.20.521329

**Authors:** Karen Doggett, Narelle Keating, Farhad Dehkhoda, Grace M. Bidgood, Evelyn Leong, Andrew Kueh, Nicos A. Nicola, Nadia J. Kershaw, Jeffrey J. Babon, Warren S. Alexander, Sandra E. Nicholson

## Abstract

Suppressor Of Cytokine Signaling (SOCS) 1 is a critical negative regulator of cytokine signaling and required to protect against an excessive inflammatory response. Genetic deletion of *Socs1* results in unrestrained cytokine signaling and neonatal lethality, characterised by an inflammatory immune infiltrate in multiple organs. Overexpression and structural studies have suggested that the SOCS1 kinase inhibitory region (KIR) and Src homology 2 (SH2) domain are important for interaction with and inhibition of the receptor-associated JAK1, JAK2 and Tyk2 tyrosine kinases, which initiate downstream signaling. To investigate the role of the KIR and SH2 domain in SOCS1 function, we independently mutated key conserved residues in each domain and analysed the impact on cytokine signaling, and the *in vivo* impact on SOCS1 function. Mutation of the SOCS1-KIR or SH2 domain had no impact on the integrity of the SOCS box complex, however, mutation within the phosphotyrosine binding pocket of the SOCS1-SH2 domain specifically disrupted SOCS1 interaction with phosphorylated JAK1. In contrast, mutation of the KIR did not affect the interaction with JAK1, but did prevent SOCS1 inhibition of JAK1 autophosphorylation. In human and mouse cell lines, both mutants impacted the ability of SOCS1 to restrain cytokine signaling, and crucially, *Socs1-R105A* and *Socs1-F59A* mice displayed a neonatal lethality and excessive inflammatory phenotype similar to SOCS1 null mice. This study defines a critical and non-redundant role for both the KIR and SH2 domain in endogenous SOCS1 function.

## Introduction

Suppressor of cytokine signaling (SOCS) proteins are critical intracellular regulators that act to limit cytokine signaling in the control of inflammation, immunity and hematopoiesis (1). The SOCS protein family consists of CIS and SOCS1-7, with CIS, SOCS1-3 primarily acting as negative feedback inhibitors of the Janus kinase-signal transducer and activator of transcription (JAK-STAT) signaling cascade. The SOCS proteins are characterised by a central Src homology 2 (SH2) domain and a C-terminal SOCS box motif (1, 2). The SOCS-SH2 domain provides specificity by binding to specific phosphorylated tyrosine residues in target proteins, and is distinguished by an extended SH2 subdomain (ESS) (3), which stabilises the phosphotyrosine binding loop (4). The SOCS box interacts with the adapters Elongin B and C, Cullin-5 and Ring Box 2 (RBX2) to form an E3 ubiquitin ligase complex, mediating target protein degradation via the proteasome (5). SOCS1 and SOCS3 are also able to directly and potently inhibit JAK catalytic activity through a unique kinase inhibitory region (KIR) and a non-canonical interaction between the SH2 domain and a GQM motif present in JAK1, JAK2 and TYK2 (3, 6). The KIR is a short amino acid stretch immediately upstream of the ESS that acts as a pseudosubstrate, blocking the substrate binding groove of JAK to prevent further kinase activity (6, 7).

Of all the SOCS proteins, SOCS1 appears to have the broadest spectrum of cytokine regulation, with expression induced by a number of cytokines, which it regulates via a classical negative feedback loop; type I and II interferons (IFN) and interleukin (IL)-2, 4, 7, 12 and 15 (8-14). This highlights the critical role SOCS1 plays in tightly regulating the biological consequences of cytokine signaling, which if left unrestrained, can generate an inappropriate immune response, inflammation and autoimmunity.

Mice with a genetic deletion of *Socs1 (Socs1-null*) are born but die before weaning with very low body weight, fatty degeneration of the liver and inflammatory infiltration of multiple organs (15). The loss of thymic cellularity and atrophy of the thymic cortex observed in these animals also highlights the important role of SOCS1 in developmental thymopoiesis (16). The critical role of *Socs1* in regulating interferon gamma (IFN*γ*) signaling is illustrated by rescue of the neonatal lethality when these mice are crossed onto an IFN*γ* null background (8) or treated with a neutralizing anti-IFN*γ* antibody (17). Genetic deletion of the *Socs1-Socs box* led to a partial loss of function, with a much less severe phenotype than observed with complete loss of SOCS1; mice showed increased responsiveness to IFN*γ*, and developed a slow and ultimately fatal inflammatory disease at around 6-months of age (18). This is consistent with the weak interaction of the SOCS1-SOCS box with Cullin-5, and implies a relatively minor role for the E3 activity of SOCS1 (19).

SOCS1 is also an important regulator of cytokine responses in humans, with SOCS1 haploinsufficiency associated with inflammatory diseases, including early onset autoimmunity, and increased STAT signaling in response to IFN*γ*, IL-2 and IL-4 (20-22).

Various *in vitro* studies have identified key residues within SOCS1 that are required for SOCS1 to regulate cytokine intracellular signaling cascades (3, 23, 24). Mutation of the conserved arginine within the SOCS1-SH2 domain (R105K/E) has been shown to disrupt binding to phosphotyrosines on JAK1, Abl, GAP and c-Kit (24-27). Similarly, overexpression and structural studies identified key residues within the KIR (Phe56, Phe59, Asp64) that are important for SOCS1 inhibitory function, with the requirement for a Phe in the KIR to be buried in a hydrophobic pocket formed by the JAK-JH1:SOCS-SH2 interaction (Phe59 in mouse SOCS1; Phe25 in mouse SOCS3), conserved in both SOCS1 and SOCS3 (3, 7, 23, 24, 28).

To investigate the *in vivo* requirement for the SH2 domain and KIR, we have generated two novel mutant mouse strains, with point mutations in either the SOCS1-SH2 domain (SOCS1-R105A) or KIR region (SOCS1-F59A) (**Figure 1A**). Both mutants displayed a similar phenotype to full *Socs1* deletion, highlighting for the first time the critical role of both the SH2 domain and KIR for SOCS1 activity *in vivo*.

**Figure 1.**
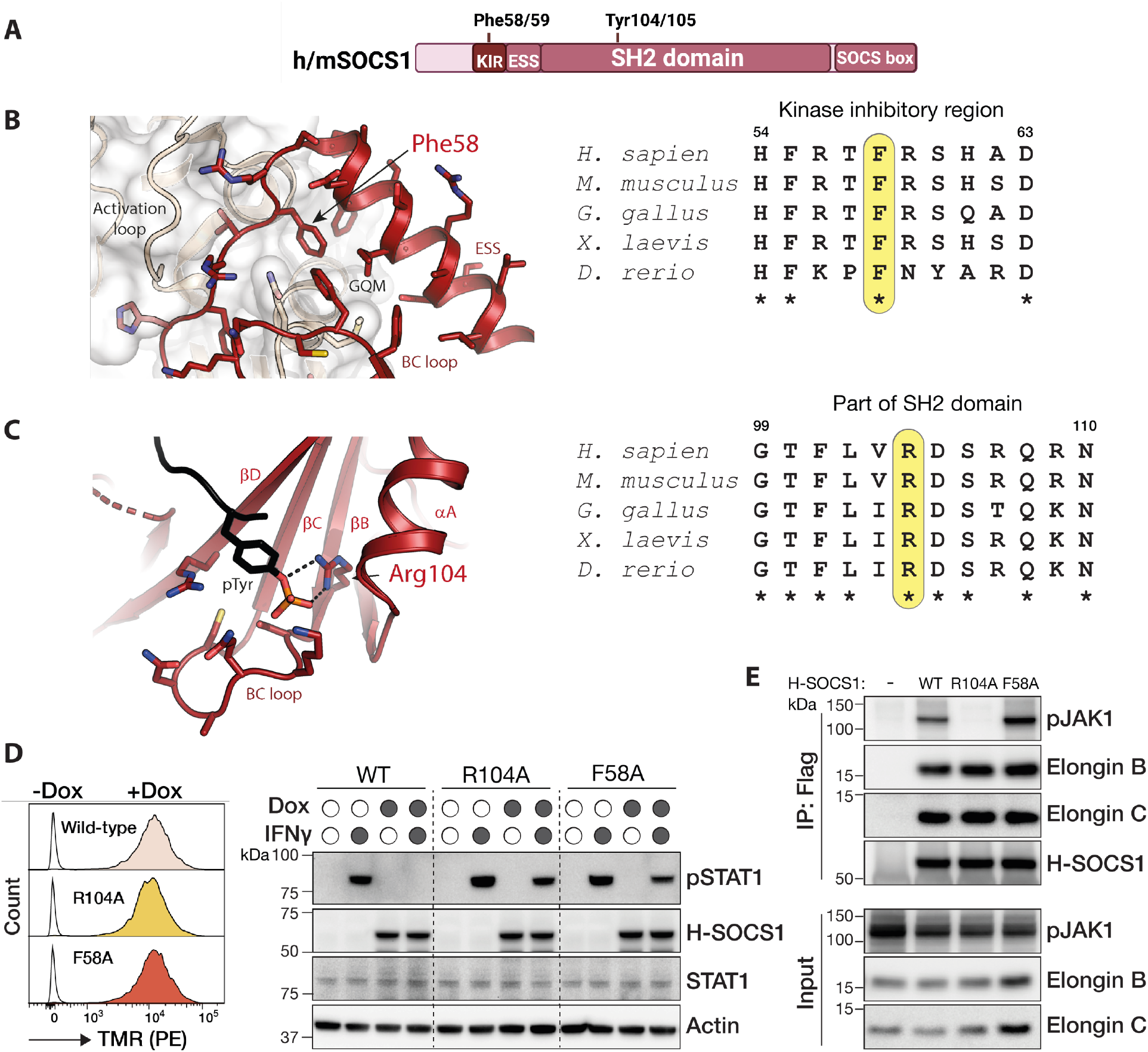
SOCS1 KIR and SH2 mutant proteins fail to negatively regulate cytokine signaling but maintain SOCS box interactions. **(A)** SOCS1 domain schematic highlighting key residues in human (h) and mouse (m) proteins. KIR: kinase inhibitory region; ESS: extended SH2 subdomain. **(B)** Crystal structure of SOCS1 in complex with the JAK kinase domain showing the interaction of *Gallus gallus* SOCS1 (red) with JAK1 (white). The conserved SOCS1 Phe residue sits within a hydrophobic pocket at the JAK1:SOCS1 interface (PDB: 6C7Y) (7). **(C)** Ribbon diagram of *Xenopus laevis* SOCS1-SH2 domain (red) complexed with a phospho-peptide derived from gp130 (black). The conserved SOCS1 Arg residue forms hydrogen-bond interactions (dotted black lines) with the phosphate on the Tyr residue (PDB: 6C5X)(7). Alignment of SOCS1 KIR (a.a. 56-67) and SH2 domain (a.a. 80-175) sequences from different species are shown to the right of the relevant structures. Multiple sequences were obtained from UniProt database and aligned using UniProt alignment tool. Mutated residues are highlighted in yellow. *indicates conserved residues. **(D)** Mutation of either Arg104 or Phe58 results in loss of SOCS1 inhibitory activity. Human A549 lung adenocarcinoma cells were treated with doxycycline (Dox) for 24 h to induce Halo-3F-hSOCS1 expression (H-SOCS1), prior to treatment with 10 ng/mL IFN*γ* for 30 min. Cells were lysed and analyzed by immunoblotting with antibodies to phosphorylated (p)STAT1, total STAT1, Halo or *β*-actin. Halo-tag expression levels detected using the fluorescent Halo-ligand tetramethylrhodamine (TMR-PE) show comparable expression of the Halo-SOCS1 wild-type (WT) and mutant (R104A, F58A) constructs. Dox-induction of *Halo-Socs1*, but not *Halo-Socs1-F58A* or *Halo-Socs1-R104A*, efficiently inhibited IFN*γ*-induced pSTAT1. **(E)** Mutation of Halo-SOCS1-R104 results in loss of JAK1 interaction but retention of the SOCS box interaction with elongins B & C. A549 cells were treated with Dox for 24 h to induce Halo-3F-SOCS1 (H-SOCS1) wild-type (WT) or mutant (R104A or F58A) expression, prior to treatment with sodium pervanadate for 15 min to inhibit phosphatase activity. Cells were lysed and Halo-3F-SOCS1 immunoprecipitated (IP) using anti-Flag beads, followed by immunoblotting for SOCS1 binding partners, pJAK1, elongin B and elongin C. Anti-Halo immunoblot showed equivalent enrichment of WT and mutant SOCS1 proteins. Immunoblotting of cell lysates (input) is shown below. *Related to Supplementary Figures 1 and 2*.

## Results

### Overexpressed SOCS1-KIR and SOCS1-SH2 mutant proteins fail to negatively regulate IFN signaling

Structural characterisation of SOCS1 bound to phosphotyrosine and the JAK1-JH1 kinase domain, confirmed the importance of Arg104 within the SH2 domain for phosphotyrosine binding, and Phe58 for KIR interaction with JAK1 (7) (**Figure 1B & C**). Although these studies were performed with xenopus and chicken SOCS1, it is obvious from sequence alignments that both regions are highly conserved (**Figure 1B & C**); strong evidence that the molecular detail of the interactions will also be conserved across species.

As further verification, we generated A549 cell lines with stable integration of doxycycline (dox)-inducible constructs encoding human *Socs1, Socs1-R104A* (SH2 mutant) or *Socs1-F58A* (KIR mutant) with N-terminal Halo and triple-Flag (3F) tags. Lentiviral transduced cells were treated overnight with doxycycline to induce Halo-SOCS1, Halo-SOCS1-R104A or Halo-SOCS1-F58A, and stimulated with 10 ng/mL IFN*γ* for 30 min, prior to analysis of the IFN*γ* signaling response by immunoblotting. Expression of Halo-SOCS1 efficiently inhibited IFN*γ*-induced phosphorylation (p) of STAT1, in contrast to Halo-SOCS1-R104A and Halo-SOCS1-F58A, which had impaired inhibitory activity, despite being expressed at comparable levels to wild-type SOCS1 (**Figure 1D**).

To investigate the impact of the point mutations on SOCS1 protein complexes we utilised the triple Flag-tag to perform immunoprecipitation experiments from A549 cell lysates. Halo-SOCS1, Halo-SOCS1-F58A and Halo-SOCS1-R104A enriched elongin B and elongin C to comparable levels, indicating the SOCS box complex was not disrupted by either the F58A or R104A mutation. Halo-SOCS1 and Halo-SOCS1-F58A, but not Halo-R104A, also enriched pJAK1 (**Figure 1E & Supplementary Figure 2**). This indicates that SOCS1-SH2 interaction with phosphotyrosine (pTyr), but not SOCS1-Phe58, is required for stable interaction with phosphorylated JAK1.

### SOCS1 KIR and SH2 point mutations result in *Socs1* neonatal lethality

To investigate the impact of mutating the KIR or SH2 domain *in vivo*, we used CRISPR/Cas9 gene editing to generate mice bearing the corresponding point mutations, either *Socs1-F59A* or *Socs1-R105A*. Correct gene targeting was obtained in C57BL/6 mice and validated by next-generation sequencing (NGS) and Sanger sequencing (**Supplementary Figure 3**).

Homozygous *Socs1*^*F59A/F59A*^ (*Socs1-F59A*) and *Socs1*^*R105A/R105A*^ (*Socs1-R105A*) mice were born in the expected mendelian ratios (**Supplementary Table 1**) but were severely runted and did not survive to weaning (3-weeks), dying between day 10-17 after birth (**Figure 2A**). The timing of this neonatal lethality matched that observed for *Socs1-null* mice (green; and previously published data) (15). Histological analysis of these pups at day 9-11 confirmed multi-organ inflammatory immune infiltration **(Figure 2B)**. Consistent with the phenotype in *Socs1-null mice*, both KIR and SH2 mutants displayed fatty degeneration of the liver **(Figure 2B**, orange arrows), an accumulation of immune infiltrate in the lungs (**Figure 2B**, blue arrows), and severe cortical atrophy of the thymus **(Figure 2B**, right panels). Together, this indicates that both the KIR and SH2 domain are required *in vivo* for SOCS1 function.

**Figure 2.**
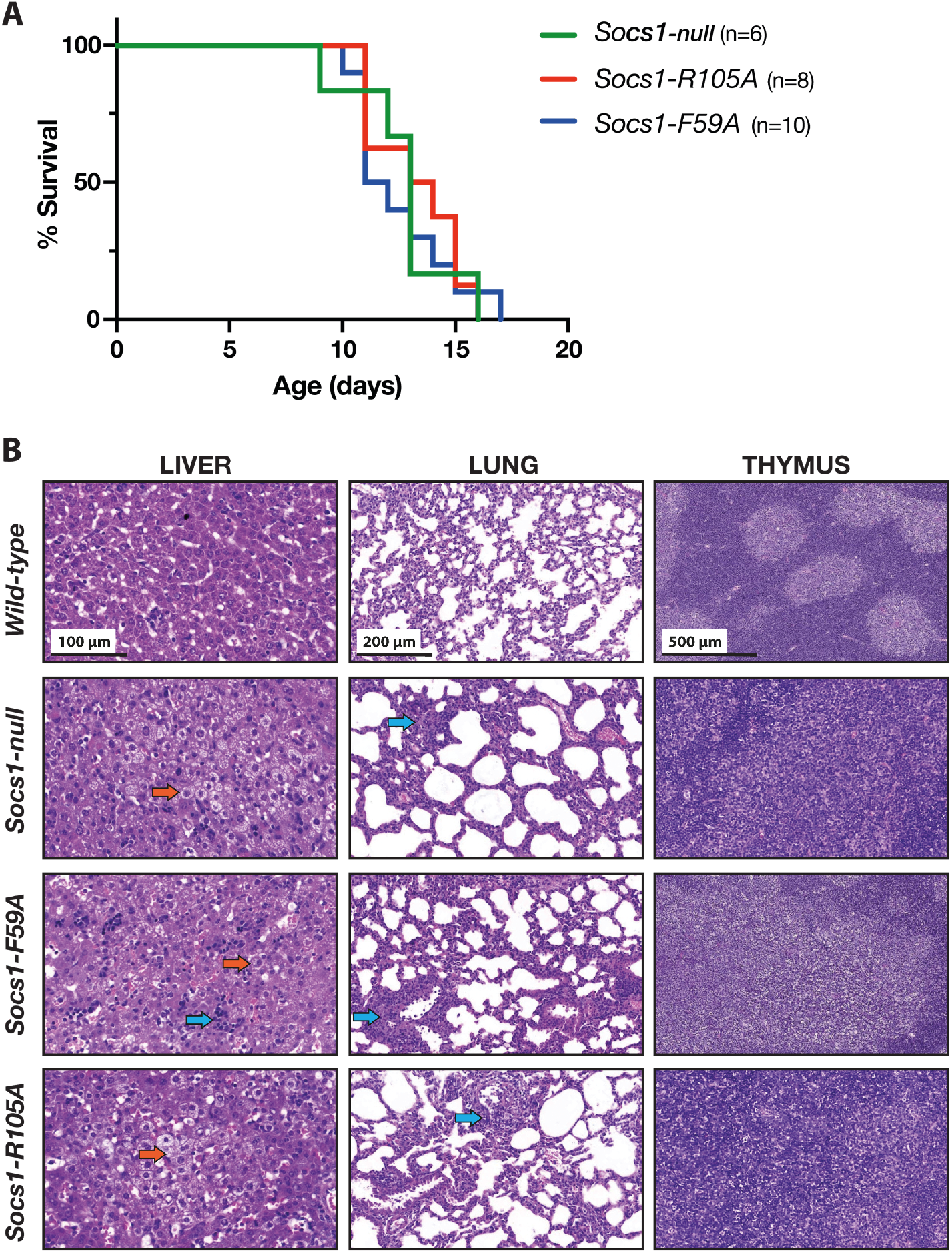
*Socs1-F59A* (KIR) and *Socs1-R105A* (*SH2)* mutants phenocopy *Socs1-null* neonatal lethality driven by global inflammation. **(A)** Kaplan-Meier survival analysis shows *Socs1-F59A* and *Socs1-R105A* mice die between 10-17 days after birth; consistent with the post-natal lethality observed in *Socs1-null* mice. n=6-10 **(B)** H&E histological analysis of neonatal tissues from *Socs1-null, Socs1-F59A* and *Socs1-R105A* mice indicates a similar degree of immune infiltrate and fatty degeneration in the liver (orange arrows), increased cellularity and macrophage accumulation in the lung (blue arrows), and atrophy of the thymus at 9-11 days of age. Images are representative of 2-4 mice per genotype.

The lethality associated with complete loss of SOCS1 can be rescued by crossing to an *Ifng* null background (8). Similarly, crossing both the *Socs1-F59A* and *Socs1-R105A* mice to *Ifng*^*-/-*^ mice rescued the neonatal lethality (data not shown), resulting in homozygous animals able to reach adulthood and reproduce, and confirming the phenotype is primarily driven by excessive IFN*γ* signaling during development.

### Mutation of either the Socs1 KIR (F59A) or SH2 domain (R105A) abrogates Socs1 inhibition of IFN signaling

The viability of adult mutant mice on an *Ifng*^*-/-*^ background enabled us to generate bone marrow-derived macrophages (BMDMs) and further characterise the effect of the mutations on cytokine signaling. Importantly, IFN*γ* and IFN*α* treatment of *Socs1-F59A Ifng*^*-/-*^ and *Socs1-R105A Ifng*^*-/-*^ BMDMs resulted in similar kinetics of SOCS1 protein induction, to that observed in wild-type control cells (*Socs1*^*+/+*^ *Ifng*^*-/-*^) (**Figure 3A & 3B**). SOCS1-R105A protein levels were comparable to wild-type SOCS1, however, we consistently observed increased levels of SOCS1-F59A (**Figure 3 and Supplementary Figure 4**).

**Figure 3.**
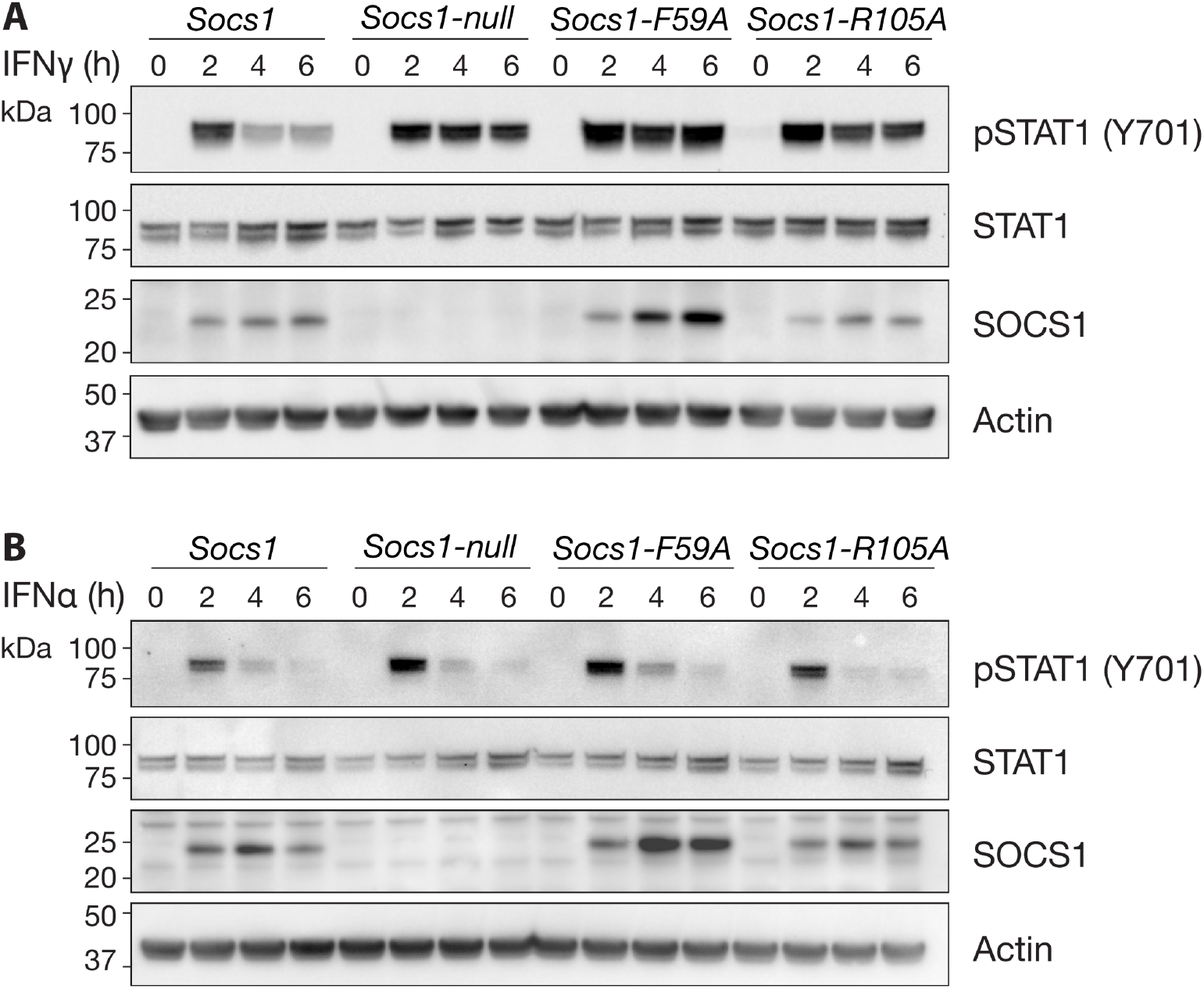
Germ-line mutation of either the *Socs1*-KIR or SH2 domain results in enhanced interferon signaling. **(A & B)** *Socs1*^*+/+*^, *Socs1-null, Socs1-F59A* (KIR mutant) and *Socs1-R105A* (SH2 mutant) *Ifng*^*-/-*^ BMDMs were treated for 0, 2, 4 or 6 h with **(A)** 100 ng/mL IFN*γ* or **(B)** 5 U/mL IFN*α*. Cells were lysed and analyzed by immunoblotting for IFN signaling responses. Representative of 3 and 2 independent experiments (A and B, respectively). *Related to Supplementary Figure 4*.

IFN*γ*-induced phosphorylation of STAT1 was enhanced in *Socs1-F59A Ifng*^*-/-*^ and *Socs1-R105A Ifng*^*-/-*^ BMDMs, in comparison to *Socs1*^*+/+*^ *Ifng*^*-/-*^ BMDMs, which displayed a drop in pSTAT1 levels at 4 and 6 h (**Figure 3A**). IFN*α*-induced pSTAT1 was also enhanced in *Socs1-F59A Ifng*^*-/-*^ and *Socs1-R105A Ifng*^*-/-*^ BMDMs, although the effect was not as pronounced as that observed with IFN*γ* stimulation (**Figure 3B and Supplementary Figure 4B**). This result is consistent with loss of negative regulation by SOCS1 and was similar to the impact of full *Socs1* deletion (**Figure 3 and Supplementary Figure 4**). These data indicated that although both mutant SOCS1 proteins were synthesized and stable *in vivo*, they were unable to negatively regulate IFN*γ* and IFN*α* signaling, consistent with data generated by overexpression of the human SOCS1 mutant proteins (**Figure 1D**). SOCS1 function was significantly impaired by both F59A and R105A, implying a critical role at the level of endogenous protein, for both the SOCS1-KIR, and SOCS1-SH2 interaction with phosphotyrosine.

## Discussion

Previous overexpression studies have suggested that both the KIR and SH2 domain are required for SOCS1 to inhibit cytokine signaling. Here we have used gene editing in the mouse to show that Phe59 in the KIR region and the phosphotyrosine binding capacity of the SH2 domain are essential for endogenous SOCS1 function. Mice bearing either the F59A or R105A mutation died shortly after weaning, consistent with full *Socs1* deletion, and displayed a similar degree of multi-organ immune infiltration and fatty degeneration of the liver, as seen in *Socs1-null* mice. Furthermore, lethality was rescued by crossing onto an *Ifng*^*-/-*^ background, indicating unrestrained IFN*γ* signaling was a major contributing cause of lethality, and again, phenocopying *Socs1* null mice.

Loss of function was not due to protein de-stabilisation, as both mutant proteins were expressed at greater or equivalent levels to wild-type SOCS1, and retained the ability to interact with the SOCS box adaptor proteins, elongins B and C. It is worth noting that mice heterozygous for *Socs1* are viable, and do not display signs of inflammatory disease until they reach 4-6 months of age (8); further evidence that the neonatal lethality observed in *Socs1-F59A* and *Socs1-R105A* mice results from loss of protein function, rather than reduced protein expression.

Interestingly, SOCS1-F59A protein was consistently induced to a greater degree than either wild-type SOCS1 or SOCS1-R105A. It is possible that F59A somehow stabilises the SOCS1 protein, so that it accumulates over time. Alternatively, given the SH2 domain retains the ability to interact with phosphotyrosine, it may act as a dominant negative, maintaining the ability to interact with phosphorylated JAK but unable to inhibit it. In this instance, bound SOCS1 may prevent de-phosphorylation of JAK by a phosphatase. Consequently, there may be an increase in JAK-mediated signaling (above *Socs1-null*) that results in greater induction of *Socs1-F59A*.

Phe59 is a key residue in the KIR and occupies a hydrophobic pocket at the interface of the SOCS1-SH2 domain and the JAK1-JH1 domain. Mutation of Phe59 in the recombinant protein abrogates the ability of SOCS1 to inhibit JAK2 (7). Somewhat surprisingly, mutation of Phe58 in the human Halo-SOCS1 construct did not disrupt the interaction with JAK1, despite complete loss of SOCS1 inhibitory activity. It is possible that mutation of Phe58/59 prevents the KIR from locking into the substrate binding site, while retaining sufficient interactions to maintain the complex. In contrast, mutation of the conserved arginine involved in binding phosphorylated tyrosine, resulted in complete loss of binding to pJAK1, and consequently, loss of inhibitory activity.

The phosphotyrosine target of the SOCS1-SH2 domain has been a subject of debate in the SOCS field. Yoshimura and colleagues proposed that the SOCS1-SH2 domain interacts with the phosphorylated JAK activation loop (24), and indeed, the SOCS1-SH2 domain binds to peptides corresponding to the phosphorylated activation loop with sub-micromolar affinity (7). However, NMR studies and examination of the full JAK-JH1 crystal structures indicate the activation loop is not accessible to the SOCS1-SH2 P0 pocket when SOCS1 is bound to the JAK-GQM motif, with the KIR occupying the substrate binding site (7). This steric hindrance could result from the activation status of recombinant JAK-JH1, with activation of JAK in cells releasing the activation loop for SOCS1-SH2:pTyr binding. Alternatively, an activated JAK dimer could enable KIR-BC loop binding to one JAK-JH1, while the SH2 domain interacts with the phosphorylated activation loop of a second JAK molecule, something that has not been captured structurally. A third possibility is that the SOCS1-SH2 domain binds to another pTyr, either on JAK or elsewhere in the protein complex.

In conclusion, this study has demonstrated the strict functional requirement for both the SOCS1 KIR and SH2 domain, and suggests they act in concert to (1) recruit SOCS1 to the JAK complex (SH2), and (2) inhibit JAK kinase activity (KIR).

## Acknowledgements

The modified pfTRE3G-Halo-3F vector was kindly provided by Dr Jonathan Bernardini (WEHI). The authors thank Natasha Blasch and Sophia Russo for mouse husbandry. J.J.B was supported by an Australian Government National Health and Medical Research Council (NHMRC) Research Fellowship (1121755). W.S.A. was supported by NHMRC Investigator and Program Grants (1173342, 1113577). G.M.B. was supported by an Australian government Research Training Program Scholarship. This work was supported under a collaborative research project with Servier and was supported in part through Victorian State Government Operational Infrastructure Support and the Australian Government NHMRC Independent Research Institutes Infrastructure Support Scheme (IRIISS).

## Methods

### Generation of SOCS1 constructs

pUC57 vector containing DNA encoding the human *Socs1* gene with flanking BamHI and NheI restriction sites at the N- and C-termini respectively, was purchased from Genscript and sub-cloned into a lentiviral Tet-On pfTRE3G expression vector (Takara Bio Ltd), modified to contain a triple Flag epitope. SOCS1-F58A and SOCS1-R104A mutations were introduced using the Quikchange Lightning Multi Site Directed Mutagenesis kit (Agilent Technologies; sequence specific primers in **Supplemental Table 3**), according to the manufacturer’s instructions. The resulting constructs encoded human SOCS1 and SOCS1 mutants with an N-terminal Halo tag followed by a triple Flag epitope (DYKDHDGDYKDHDIDYKDDDDK) (Halo-3F-SOCS1). All constructs were confirmed by Sanger sequencing (Australian Genome Research Facility; AGRF).

### Generation of Halo-tagged human SOCS1 cell lines

Human lung adenocarcinoma (A549 RRID:CVCL_0023) and kidney epithelial (HEK293T RRID:CVCL_0063) cells were cultured in Dulbecco’s Modified Eagle Medium (DMEM; Thermo Fisher) + 10% fetal bovine serum (FBS; Thermo Fisher) at 37°C in a humidified incubator with 10% CO_2_. Cells were routinely tested for mycoplasma contamination.

HEK293T cells were transfected with lentiviral constructs (10 μg) and packaging vectors (7.5 μg of psPAX and 3 μg of VSVg) using Lipofectamine 2000 (Invitrogen) according to the manufacturer’s instructions. Media was replaced 24 h post-transfection and cells were incubated for a further 48 h. Lentivirus was harvested and passed through a 0.45 μm filter and A549 target cells were spin-infected in the presence of polybrene (8 μg/mL; 500 *g* at 37 °C for 2 h). Lentivirus was incubated with target cells for 24 h prior to recovery in fresh media for 3 days. Cells were selected in puromycin (2.5 µg/mL; Merck 58-58-2) for 7 days.

### TMR-ligand flow cytometry

Halo-3F-SOCS1 expression was induced with doxycycline overnight (1 μg/mL) and Halo-tagged protein detected by incubation with fluorescent Halo TMR ligand (10 nM; Promega) overnight. Cells were incubated with fixable viability stain 510 (FVS510) (BD Horizon; 0.38 µg/mL) for 10 min at room temperature to exclude dead cells then washed three times in phosphate-buffered saline (PBS) + 2% FCS, prior to data collection on a BD LSRFortessa. Data analysis was performed using FlowJo software v10, gating on single, viable cells.

### Anti-Flag immunoprecipitation (IP)

A549 cells containing doxycycline-inducible Halo-3F-SOCS1 constructs (10^7^ cells/IP) were treated as indicated and lysed on ice in 1 mL of lysis buffer (1% v/v NP-40, 50 mM HEPES, pH 7.4, 150 mM NaCl, 1 mM EDTA, 10% glycerol), supplemented with protease inhibitor cocktail (Calbiochem), 1 mM phenylmethylsulphonyl fluoride (PMSF), 5 mM NaF and 1 mM Na_3_VO_4_]. Cell lysates were pre-cleared using Protein A-Sepharose at 4 °C on a rotating wheel for 2-4 h. Pre-clearing beads were pelleted twice by centrifugation (15,000 *g* for 2 min at 4 °C) and discarded. Anti-Flag M2-conjugated beads (15 μL packed bead volume; Sigma) were rotated with cleared cell lysates for 3 h at 4°C. Antibody complexes were collected by centrifugation (15,000 *g* for 30 sec at 4 °C), washed 3 × 1 mL in ice cold lysis buffer and mixed with equal volume of 2x reducing sample buffer [62.5 mM Tris-HCl, pH 6.8, 10% v/v glycerol, 2% w/v SDS, 0.0025% w/v bromophenol blue, 50 mM w/v dithiothreitol (DTT)].

### Immunoblotting

Cell lysates or immunoprecipitated complexes were separated by sodium dodecyl sulphate-polyacrylamide gel electrophoresis (SDS-PAGE) and electrophoretically transferred to nitrocellulose (Amersham) or polyvinylidene fluoride PVDF (Immobilon) membranes. Membranes were blocked in 10% w/v skimmed milk (or 3% BSA w/v tris-buffered saline with 0.1% v/v Tween 20 for anti-phospho-JAK1) and incubated with primary antibody at 4°C overnight.

Anti-phospho-STAT1 (CST#7649; 1:1000) and anti-STAT1 (9172; 1:1000) antibodies were purchased from Cell Signaling Technology. Anti-phospho-JAK1 (7000028; 1:1000) antibody was purchased from Invitrogen. Anti-elongin B (ab154854; 1:2000) and anti-SOCS1 (ab9870; 1:1000) antibodies were purchased from Abcam. Anti-elongin C antibody (Clone 56/SIIIp15; 1:1000) was from BD Transduction Laboratories. Anti-Halo antibody was purchased from Promega (G9211; 1:1000). Anti-actin-HRP antibody (C4) was obtained from Santa Cruz (sc-47778 HRP; 1:1000). Antibody binding was visualized with peroxidase-conjugated sheep anti-rabbit immunoglobulin (Southern Biotech; 4010-05; 1:15000), sheep anti-mouse immunoglobulin (GE Healthcare; NA931-1ML; 1:10000) or donkey anti-goat immunoglobulin (Invitrogen; A16005) and the enhanced chemiluminescence (ECL) system (Amersham or Millipore).

### Mice

Mice carrying a germline mutation of Arg105 to Ala (*Socs1-R105A*) or Phe59 to Ala (*Socs1-F59A*) in SOCS1 were generated on a C57BL/6 J background by the MAGEC laboratory at the Walter & Eliza Hall Institute, as previously described (29). 20 ng/μL Cas9 mRNA, 10 ng/μL single guide RNAs, and 40 ng/μL donor template were injected into fertilized one-cell stage embryos. Guides and targeting vector sequences are provided in **Supplementary Table 2**. Founder mice were analyzed by next-generation sequencing (NGS) to confirm the correct sequence change. Sanger sequencing is shown in **Supplementary Figure 3**. Mice carrying the mutation were backcrossed to C57BL/6J mice for three generations to eliminate potential off-target events, and next-generation sequencing repeated. Heterozygous mice were intercrossed to generate homozygotes, and litters were tattooed and sampled for genotyping at birth.

Mice were genotyped by NGS using genomic DNA extracted from ear biopsies with the Direct PCR Lysis tail reagent (Viagen) and 5 mg/mL proteinase K (Worthington), according to the manufacturer’s instructions. Genotyping primers are shown in **Supplementary Table 3**. *Socs1-null* (15) and *Ifng*^*-/-*^ (30) mice have been described previously. Experiments were performed at the Walter and Eliza Hall Institute (WEHI) in accordance with the NHMRC Australian code for the care and use of animals for scientific purposes. All experiments were approved by the WEHI Animal Ethics Committee (AEC 2021.002).

### Histology

Tissue samples were taken at ethical endpoint from 9-11 day old *Socs1* wild-type littermate, *Socs1-null, Socs1-R105A* and *Socs1-F59A* mice, fixed with 10% buffered formalin, embedded in paraffin and stained with hematoxylin and eosin using standard techniques. Images were captured using the Panoramic Scan II (3D HISTECH Ltd.).

### Mouse bone marrow-derived macrophages (BMDMs) and IFN stimulation

Bone marrow was extracted from the femurs and tibias of 8-10-week old mice and cultured in DMEM supplemented with 15% v/v L929 cell conditioned media as a source of macrophage-colony stimulating factor (M-CSF), 100 U/mL penicillin, 0.1 ng/mL streptomycin, and 10% v/v FBS, at 37°C in a humidified incubator with 10% CO_2_. BMDMs were supplemented every second day with L929 cell conditioned media and plated for cytokine time courses on day 6. BMDMs were treated with 100 ng/mL mouse IFN*γ* or 5 U/mL IFN*α* (R&D Systems) for 2, 4 or 6 h and lysed directly into Laemmli sample buffer.

### Statistical analysis

Statistical analyses were performed with Prism 9 software. Kaplan Meier curves were not significantly different as analysed by Log-rank (Mantel-Cox) test.

