## Supplemental data for "The SOCS1 KIR and SH2 domain are both required for suppression of cytokine signaling *in vivo*"

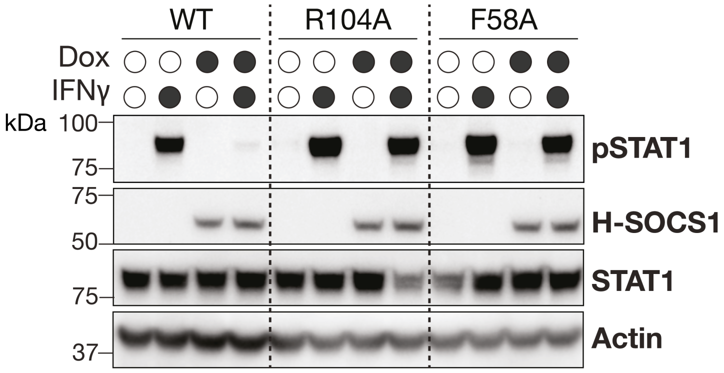


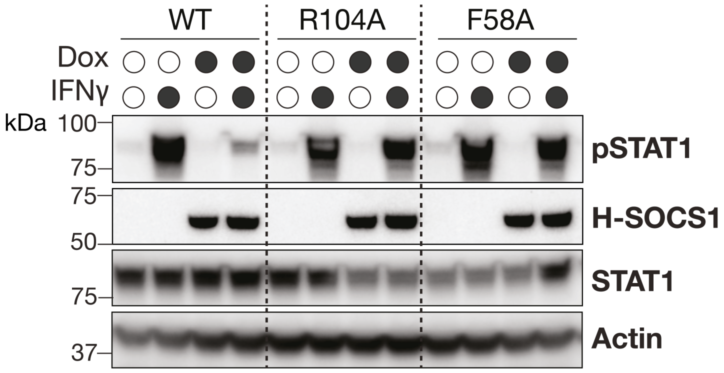


**Supplementary Figure 1. Mutation of either Arg104 or Phe58 results in loss of SOCS1 inhibitory activity.** Human A549 lung adenocarcinoma cells were treated with doxycycline (Dox) for 24 h to induce Halo-3F-human SOCS1 expression (H-SOCS1), prior to treatment with 100 ng/mL IFNγ for 30 min. Cells were lysed and analyzed by immunoblotting with antibodies to phosphorylated (p)STAT1, total STAT1, Halo or β-actin. Anti-Halo immunoblots showed comparable expression of the Halo-3F-SOCS1 wild-type (WT) and mutant (R104A, F58A) constructs. Dox-induction of *Halo-Socs1,* but not *Halo-Socs1-F58A* or *Halo-Socs1-R104A,* efficiently inhibited IFNγ-induced pSTAT1. *Experimental repeats related to Figure 1D.*

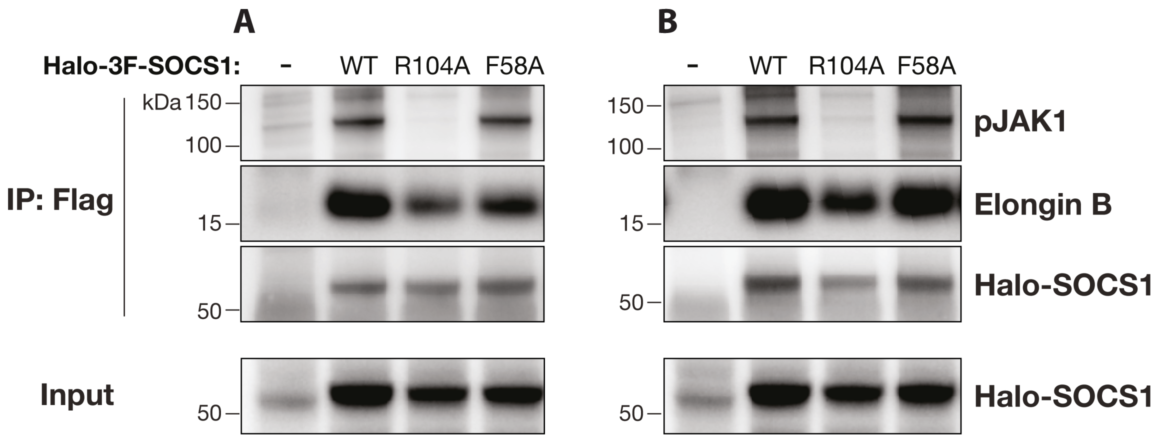


**Supplementary Figure 2. Mutation of Halo-SOCS1-R104 results in loss of interaction with JAK1 but retains the interaction with elongin B.** (**A & B**) A549 cells were treated with Dox for 24 h to induce Halo-3F-SOCS1 (H-SOCS1) wild-type (WT) or mutant (R104A or F58A) expression, prior to treatment with sodium pervanadate for 15 min to inhibit phosphatase activity. Cells were lysed and Halo-3F-SOCS1 immunoprecipitated (IP) using anti-Flag beads, followed by immunoblotting for SOCS1 binding partners, phosphorylated (p)JAK1 and elongin B. Anti-Halo immunoblot showed enrichment of WT and mutant SOCS1 proteins. Immunoblotting of cell lysates (input) for Halo-SOCS1 is shown below. Experiments in (**A**) and (**B**) were performed on different days and analyzed together by immunoblotting on the same gels. *Experimental repeats related to Figure 1E.*

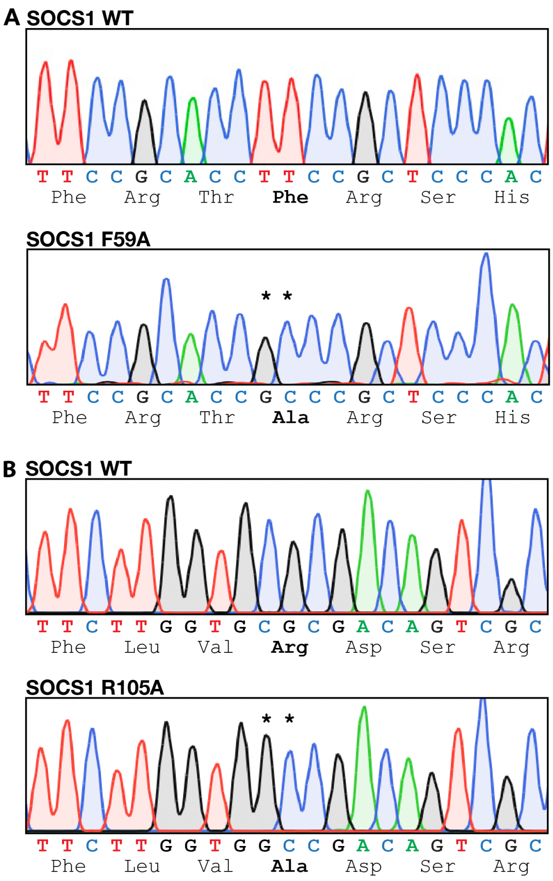


**Supplementary Figure 3. Confirmation of targeted *Socs1-F59A* and *Socs1-R105A* point mutants.** Sanger sequencing chromatograms confirming **(A)** the mouse *Socs1-F59A* and **(B)** *Socs1-R105A* sequence (homozygous mutant compared to corresponding WT sequence). *Highlights the CRISPR targeted specific base residue changes, with the amino acid change in bold.

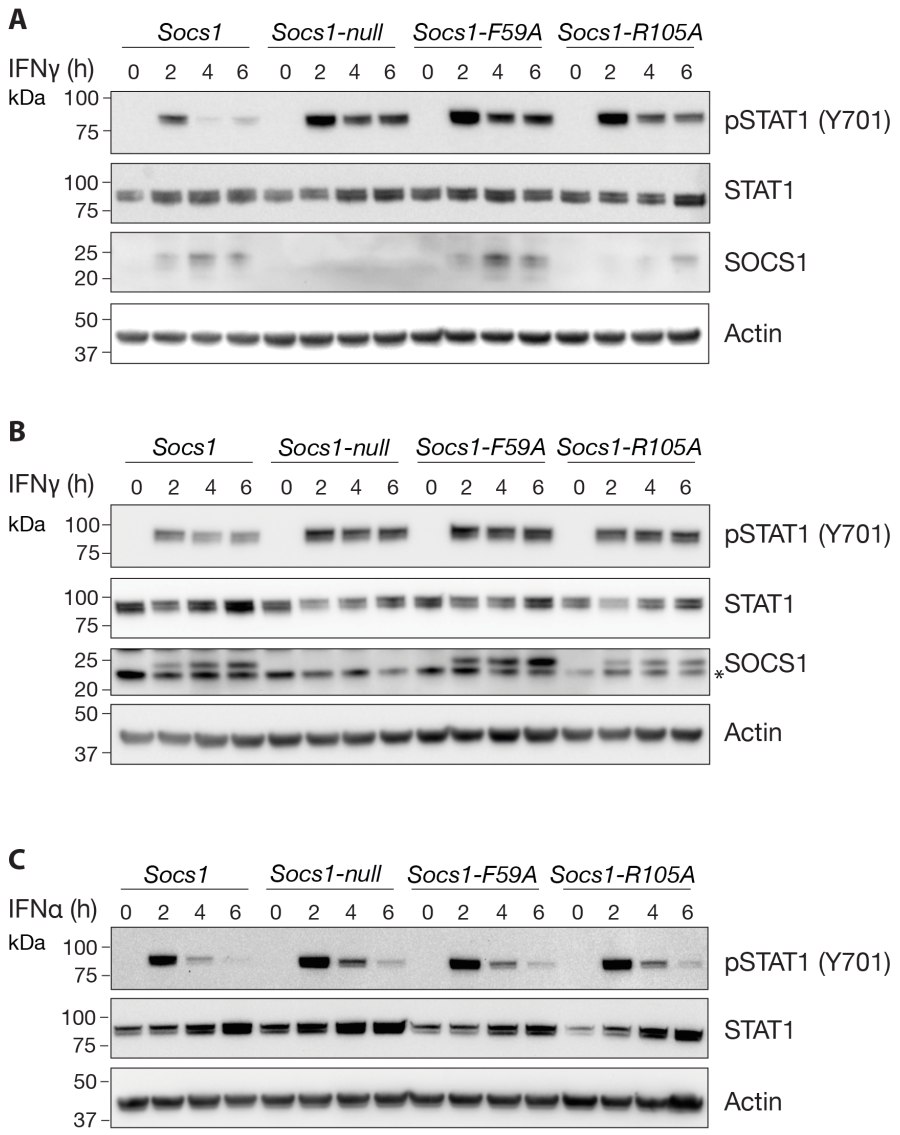


**Supplementary Figure 4. Germ-line mutation of either the *Socs1*-KIR or SH2 domain results in enhanced IFN signaling.** *Socs1*$^{+/+}$, *Socs1-null, Socs1-F59A* (KIR mutant) and *Socs1-R105A* (SH2 mutant) *Ifng*$^{-/-}$ BMDMs were treated for 0, 2, 4 or 6 h with **(A & B)** 100 ng/mL IFNγ, or (**C**) 5 U/mL IFNα. Cells were lysed and analysed by immunoblotting for IFN signaling responses. Antibodies used correspond to the labels to the right of each blot. p=phosphorylated. * indicates non-specific band. *Two additional independent experiments related to Figure 3*.

**Supplementary Table 1. Homozygous *Socs1-F59A* and *Socs1-R105A* mice were born in the expected mendelian ratios.**

**Offspring statistics from *Socs1-F59A/+* x *Socs1-F59A/+***

| | Total | +/+ | F59A/+ | F59A/F59A |
|---|---|---|---|---|
| Genotypes at **birth** | 57 | 15 | 32 | 10 |
| Predicted % | 100 | 25 | 50 | 25 |
| Observed % | 100 | 26 | 56 | 18 |

**Offspring statistics from *Socs1-R105A/+* x *Socs1-R105A/+***

| | Total | +/+ | R105A/+ | R105A/F59A |
|---|---|---|---|---|
| Genotypes at **birth** | 26 | 8 | 12 | 6 |
| Predicted % | 100 | 25 | 50 | 25 |
| Observed % | 100 | 31 | 46 | 23 |

**Supplementary Table 2. Guide and targeting sequences.**

### Single guide (sg) sequences. Used for generation of KIR (F59A) and SH2 (R105A) mutants

sgRNA1: acggcgccccgctcgagcga

sgRNA2: ggctagttgtggtttatgac

### Targeting vector sequences

**F59A:**

taggaattgaactcaggacctctggtagagccgtcagtgcttttaaccgcccggccatctctccagcctgagtcagcagttttaattttttaaataaagcattcctatccaca
agggatgcttgcacgcaagtgtacacagcagacatgacagcaaggcaacattgcttggctaacggaccaagttcagggaactgaacttcccagagctttcccagaa
gtgggacaggctatctacctcaccgcacaatctcccagtccctccctctcccatctgcgttgacactttcaaaatcacacaggggcagtgggcagtacttcccagcagg
acactgtgaatgctggaaaaaggtggcttgcaggctaggctatggaaccacgttttgcatccaagcctgcaggaccctacaatggcagccactgtacagtgggtactg
ggctctctggacccctgggggtgggtcaggaagcagctatacagctcagcatgtttctttttcacaacttttagaaactgaaaccaagttgtccaaagacagcttgacttg
ccctgtactttagatggaaggaaggcaatcagatccctggcatatatactaaagactgcatcactggaatgtgatggatgggagagggctagccaggggcagggcg
agaggacacaccctctgactgaggaggaggggagatttcctgagagcaggcagaggtggaggcctggagtcctctatttatagggttcttctgaaaaggtctgccaa
gagcttggcagggaggcaggcctgcaggagccagcccagccttgtcacccacctcagcaccaataagcctgccaagtcccaaggggatactgtaccccaagag
gttttcctcactttgcctgggctggaagcacagtttggggatggagttcaccaaccccagtactactaaaataggctggtacaaggccctgtgtttagtccccagcactgc
aaacaagcatccgtgatgcttgccagaccagaaagacctgaaggccagtctaggctgagacgcgctagagggaggcagtgctacaaacccagcctttatcgccc
accaagggttagactgaccatggctgcaccacaaccacagccacacaaattcaagcaccacaagataagtttggttgtgcttaggatgaaatatggcgctctgtgctc
tgtctggggagaccaggacttggcctcagagcttcagaggagccgcctgacagctaacagagaaacctgtagagggcagtgtgcgtcactgtgagcacagggaa
gcctgcacgcgaacctcatccgccttcattcataaacatcgtcagccaggcacccactcctggcctttcaggacaaactgaatcacgaaaccacacagtgtcctaatg
ataggtctgactgcctggatcctaagccaaggtgtgctcggggccccttgtacaggcagatgcttgctggagtcttgaccaaaagtttggggattacagcttttttggaat
cacagccccgcccggggcctcagtttctccggctgcccacgtagtaagagtgcagagagtgagtccaggcccctgggaacccagcccaaccccgcccggtttccg
aggaactaggccgggagcgggggcgcccctcccgcaccgccttaggcttcctttgaagcctctgcggtcaggccaccgcttcctgggaagcccaagccaaggcca
ggccgaatggccaacgggaggggcccgcacgcggctctggaggagggcggcggccccacaggtctctaggactagctagccgggttccaagaagggtcgaga
ttgccaaggccttcgggtcctgggcaggaaggaccctggcagggaggagctgcttggggagcacagggtccaggcgaggcggagccctaaccagaagaatgca
gacacccggagggaggaggcgtgtcagccccgcgctagcatcccacacggcgggctgcgatttggggcgaaggtagagcaaaagagcgggcaccaagtcct
aagcaccccaggcacggaacacaagattccggttggagccggaaccccagaggtcccaatgtgggaaggtgcgaggcgaaaccaagttagaggaaccgtctc
cagggagagcctcaggagtctagagagaacccgaaagacttgccggaaagagaaaccgaaagcgggggtgggctggacctgtgggcggggcctgcctggttta
agagcctgatgcaggggcgggcagcAGCAGAGAGAACTGCGGCCGTGGCAGCGGCACGGCTCCCAGCCCCGGAGCATGC
GCGACAGCCGCCCCGGAGCCCCCAGCCGCGGCTCCCCGCGTCCTGCCGCCAGgtgagccaaggcagctgcgagggagca
ggcgggaggggatgggaggtgatgggagcagagcccggcaggactaacctcgcagactgtatggcagggtcgaggatgccacgcctctggcgccccccaccc
ccgccccggctcccccaggaggcTTctcgctcgagcggggcgccgtcagcccctcctctccggccctgagccctgatcctccgcccgggttTTagtccccggcgtg
gccagtaggcggcagccgcgaggcgactagccaccagcggggacggccgggagtcgggcccctctccacgccccсttctccacgcgcgcggggaggcagggc
tccaccgccagtctggaagggttccgcatacaggaacggcctacttcgcagATGAGCCCACCGAGGCTCAAGCTCCGGGCGGATTCTGC
GTGCCGCTCTCGCTCCTTGGGGTCTGTTGGCCGGCCTGTGCCACCCGGACGCCCGGCTCACTGCCTCTGTCTC
CCCCATCAGCGCAGCCCCGGACGCTATGGCCCACCCCTCCAGCTGGCCCCTCGAGTAGGATGGTAGCACGCAA
CCAGGTGGCAGCCGACAATGCGATCTCCCCGGCAGCAGAGCCCCGACGGCGGTCAGAGCCCTCCTCGTCCTC
GTCTTCGTCCTCGCCAGCGGCCCCCGTGCGTCCCCGGCCCTGCCCGGCGGTCCCAGCCCCAGCCCCTGGCGA
CACTCACTTCCGCACCGCCCGCTCCCACTCCGATTACCGGCGCATCACGCGGACCAGCGCGCTCCTGGACGCC
TGCGGCTTCTATTGGGGACCCCTGAGCGTGCACGGGGCGCACGAGCGGCTGCGTGCCGAGCCCGTGGGCACC
TTCTTGGTGCGCGACAGTCGCCAACGGAACTGCTTCTTCGCGCTCAGCGTGAAGATGGCTTCGGGCCCCACGA
GCATCCGCGTGCACTTCCAGGCCGGCCGCTTCCACTTGGACGGCAGCCGCGAGACCTTCGACTGCCTTTTCGA
GCTGCTGGAGCACTACGTGGCGGCGCCGCGCCGCATGTTGGGGGCCCCGCTGCGCCAGCGCCGCGTGCGGC
CGCTGCAGGAGCTGTGTCGCCAGCGCATCGTGGCCGCCGTGGGTCGCGAGAACCTGGCGCGCATCCCTCTTAA
CCCGGTACTCCGTGACTACCTGAGTTCCTTCCCCTTCCAGATCTGACCGGCTGCCGCTGTGCCGCAGCATTAAG
TGGGGGCGCCTTATTATTTCTTATTATTAATTATTATTATTTTTCTGGAACCACGTGGGAGCCCTCCCCGCCTGGG
TCGGAGGGAGTGGTTGTGGAGGGTGAGATGCCTCCCACTTCTGGCTGGAGACCTCATCCCACCTCTCAGGGGT
GGGGGTGCTCCCCTCCTGGTGCTCCCTCCGGGTCCCCCCTGGTTGTAGCAGCTTGTGTCTGGGGCCAGGACCT
GAATTCCACTCCTACCTCTCCATGTTTACATATTCCCAGTATCTTTGCACAAACCAGGGGTCGGGGAGGGTCTCT
GGCTTCATTTTTCTGCTGTGCAGAATATCCTATTTTATATTTTTACAGCCAGTTTAGGTAATAAACTTTATTATGAAA
GTTTTTTTTTAAAAGAAACAAagattcctagagcgtatgctttggccaaacgtcctgggttgggagtggggtatacagactgacttttcttgaagtcttcggga
tgctgggggagggggggagggtcggacatcatatacatctccacccacagtgatggggaccaaacttccaggctagttgtggtttatgactTTgaagatggccgctc
ctgagtatccgtgcctggtctctggtatttctgtgatgggatcctacagggacagcccctgacactgagtactgtgttgcccccagtatacagaggagaaaactgagaga
cgggtaattgacgacagaccattcctggactggagaggtgggccttttaactgtccatcctgcatcaatttgaaatggatgacagagaggaaacttctttgcttctctgac
cacaactacttccaggaagagggtggggcagagaggccagggctccccgagtttttTTaaggagtggtggttctgcaagccatcaagcatttcagagggatgatgg
gccatgctggcttgacttttccctcctctccccactctgccttgtgtactttttggagccagaaagctgctttgtggtctgtggtggcaggggcctgggtgctgctggtggcttg

ggcaattccctttgcctctgggcctctgaagcaatggtgctgttaggagcccagctggaagggctgttccaagacacaagcagggcactagaaaggacccagcact
ggctctgagacagacgcctggagcttgctggtcatcaggggcccacatcccacaccatcttcatgagttctgtgactcctgcagcctgggacatgcttctagtggctttcct
aatgcttaatgatcacgttcagaggaagtggctgctctgggcacactgacactgtctgggctttctctggagcgttagaatggctgtggttctccatcccagcatggccagt
ccttcagggaggtgaggagagagcatggaggaccaaacctgctgtgtgaatgaataaggttctccccatgctgcttttcagtctagcagggcgaagggcagagacct
ggtgagtactgtggtaccatgcagatccccacagcaagagcttgggggtggagggaggaggcagctagctgggaaacccctgcttgacacctggacccgccagg
gagttaagcccctgcctttggggaacaggctggccagaaggcaaagaaaagctcttggggtgaggcctcaccgttgcctggcttgtactggtactcaccagttccagg
tcacctttaaaacagttgcccttcacctgggcttctggttggccctgtggcactgctggcatccctgtttcacaggctggaaccaggccctccttcagacatggtgaataga
gaagctgttcttcagcatgccagggggatttcctatctgaagttgacaggtcctgttactcacacatctaaatcacagatccgctgttctgtaaaagcctgaggtagaaca
gctaggaataaaggctcccgtcacactccctggagctggccccttcttccctttgtgagtcattaaaaaaaaaaaaaaaaaaaaaatccatcctgacatatggaaggtg
acctaaaagcccctgggagctacacttcagtgtctgtgtgtctgtggccctgtggcttcccgggggtacagattctctatctggcctcaaacctcagcctcatgattcagca
gagttgactgacgtcagcgcacaagaatgaaaagtgagaggagggctcaagtagggggccaggtggggtgtggagctggccgactgggaaaggaacagggct
gcttgcacacgcttagaagccgccgcagctggcaactgtggtccttgttttactgaggaggaagaggctcaggccttgaaggaggcctgacctgttttcagcaagatga
atttattattttttttcttttggtttttgagacagggttcctctgtgtatccccagctatcctggaactcactctgtagaccaggctgcctctgcctggtgcacctctggagtgcacta
ccactcctgtgaaagtacattgctaggctcccctcaccccgtaaactctcagcctagggagtttagttaggtgtgggatgggcaggcgagcctcttctatggggtgtccag
cacaaagcacttagcacagagacttgatacagtagaggcaacaggtacacaatgaaggacgtgcagagagttcaaggtctgcctgggctacatagtgagaaata
agaaacactgcctgtttccgatgttcttctaagccccaagaacacagaaataaatggaaataaatattctcttggaggataacagacaaatataaaggaacactcctct
ttaagaagaggcagtgctcagaggaagactaatggtggttggatgtgggcagcactgga

**R105A:**

taggaattgaactcaggacctctggtagagccgtcagtgcttttaaccgcccggccatctctccagcctgagtcagcagttttaattttttaaataaagcattcctatccaca
agggatgcttgcacgcaagtgtacacagcagacatgacagcaaggcaacattgcttggctaacggaccaagttcagggaactgaacttcccagagctttcccagaa
gtgggacaggctatctacctcaccgcacaatctcccagtccctccctctcccatctgcgttgacactttcaaaatcacacaggggcagtgggcagtacttcccagcagg
acactgtgaatgctggaaaaaggtggcttgcaggctaggctatggaaccacgttttgcatccaagcctgcaggaccctacaatggcagccactgtacagtgggtactg
ggctctctggacccctgggggtgggtcaggaagcagctatacagctcagcatgtttctttttcacaacttttagaaactgaaaccaagttgtccaaagacagcttgacttg
ccctgtactttagatggaaggaaggcaatcagatccctggcatatatactaaagactgcatcactggaatgtgatggatgggagagggctagccaggggcagggcg
agaggacacaccctctgactgaggaggaggggagatttcctgagagcaggcagaggtggaggcctggagtcctctatttatagggttcttctgaaaaggtctgccaa
gagcttggcagggaggcaggcctgcaggagccagcccagccttgtcacccacctcagcaccaataagcctgccaagtcccaaggggatactgtaccccaagag
gttttcctcactttgcctgggctggaagcacagtttggggatggagttcaccaaccccagtactactaaaataggctggtacaaggccctgtgtttagtccccagcactgc
aaacaagcatccgtgatgcttgccagaccagaaagacctgaaggccagtctaggctgagacgcgctagagggaggcagtgctacaaacccagcctttatcgccc
accaagggttagactgaccatggctgcaccacaaccacagccacacaaattcaagcaccacaagataagtttggttgtgcttaggatgaaatatggcgctctgtgctc
tgtctggggagaccaggacttggcctcagagcttcagaggagccgcctgacagctaacagagaaacctgtagagggcagtgtgcgtcactgtgagcacagggaa
gcctgcacgcgaacctcatccgccttcattcataaacatcgtcagccaggcacccactcctggcctttcaggacaaactgaatcacgaaaccacacagtgtcctaatg
ataggtctgactgcctggatcctaagccaaggtgtgctcggggccccttgtacaggcagatgcttgctggagtcttgaccaaaagtttggggattacagcttttttggaat
cacagccccgcccggggcctcagtttctccggctgcccacgtagtaagagtgcagagagtgagtccaggcccctgggaacccagcccaaccccgcccggtttccg
aggaactaggccgggagcgggggcgcccctcccgcaccgccttaggcttcctttgaagcctctgcggtcaggccaccgcttcctgggaagcccaagccaaggcca
ggccgaatggccaacgggaggggcccgcacgcggctctggaggagggcggcggccccacaggtctctaggactagctagccgggttccaagaagggtcgaga
ttgccaaggccttcgggtcctgggcaggaaggaccctggcagggaggagctgcttggggagcacagggtccaggcgaggcggagccctaaccagaagaatgca
gacacccggaggggaggaggcgtgtcagccccgcgctagcatcccacacggcgggctgcgatttggggcgaaggtagagcaaaagagcgggcaccaagtcct
aagcaccccaggcacggaacacaagattccggttggagccggaaccccagaggtcccaatgtgggaaggtgcgaggcgaaaccaagttagaggaaccgtctc
cagggagagcctcaggagtctagagagaacccgaaagacttgccggaaagagaaaccgaaagcgggggtgggctggacctgtgggcggggcctgcctggttta
agagcctgatgcaggggcgggcagcAGCAGAGAGAACTGCGGCCGTGGCAGCGGCACGGCTCCCAGCCCCGGAGCATGC
GCGACAGCCGCCCCGGAGCCCCCAGCCGCGGCTCCCCGCGTCCTGCCGCCAGgtgagccaaggcagctgcgagggagca
ggcgggaggggatgggaggtgatgggagcagagcccggcaggactaacctcgcagactgtatggcagggtcgaggatgccacgcctctggcgccccccaccc
ccgccccggctcccccaggaggcTTctcgctcgagcggggcgccgtcagcccctcctctccggccctgagccctgatcctccgcccgggttTTagtccccggcgtg
gccagtaggcggcagccgcgaggcgactagccaccagcggggacggccgggagtcgggcccctctccacgccccttctccacgcgcgcggggaggcagggc
tccaccgccagtctggaagggttccgcatacaggaacggcctacttcgcagATGAGCCCACCGAGGCTCAAGCTCCGGGCGGATTCTGC
GTGCCGCTCTCGCTCCTTGGGGTCTGTTGGCCGGCCTGTGCCACCCGGACGCCCGGCTCACTGCCTCTGTCTC
CCCCATCAGCGCAGCCCCGGACGCTATGGCCCACCCCTCCAGCTGGCCCCTCGAGTAGGATGGTAGCACGCAA
CCAGGTGGCAGCCGACAATGCGATCTCCCCGGCAGCAGAGCCCCGACGGCGGTCAGAGCCCTCCTCGTCCTC
GTCTTCGTCCTCGCCAGCGGCCCCCGTGCGTCCCCGGCCCTGCCCGGCGGTCCCAGCCCCAGCCCCTGGCGA
CACTCACTTCCGCACCTTCCGCTCCCACTCCGATTACCGGCGCATCACGCGGACCAGCGCGCTCCTGGACGCCT
GCGGCTTCTATTGGGGACCCCTGAGCGTGCACGGGGCGCACGAGCGGCTGCGTGCCGAGCCCGTGGGCACCT
TCTTGGTGGCCGACAGTCGCCAACGGAACTGCTTCTTCGCGCTCAGCGTGAAGATGGCTTCGGGCCCCACGAG
CATCCGCGTGCACTTCCAGGCCGGCCGCTTCCACTTGGACGGCAGCCGCGAGACCTTCGACTGCCTTTTCGAG
CTGCTGGAGCACTACGTGGCGGCGCCGCGCCGCATGTTGGGGGCCCCGCTGCGCCAGCGCCGCGTGCGGCC
GCTGCAGGAGCTGTGTCGCCAGCGCATCGTGGCCGCCGTGGGTCGCGAGAACCTGGCGCGCATCCCTCTTAAC
CCGGTACTCCGTGACTACCTGAGTTCCTTCCCCTTCCAGATCTGACCGGCTGCCGCTGTGCCGCAGCATTAAGT
GGGGGCGCCTTATTATTTCTTATTATTAATTATTATTATTTTTCTGGAACCACGTGGGAGCCCTCCCCGCCTGGGT
CGGAGGGAGTGGTTGTGGAGGGTGAGATGCCTCCCACTTCTGGCTGGAGACCTCATCCCACCTCTCAGGGGTG
GGGGTGCTCCCCTCCTGGTGCTCCCTCCGGGTCCCCCCTGGTTGTAGCAGCTTGTGTCTGGGGCCAGGACCTG
AATTCCACTCCTACCTCTCCATGTTTACATATTCCCAGTATCTTTGCACAAACCAGGGGTCGGGGAGGGTCTCTG
GCTTCATTTTTCTGCTGTGCAGAATATCCTATTTTATATTTTTACAGCCAGTTTAGGTAATAAACTTTATTATGAAAG
TTTTTTTTTAAAAGAAACAAagattcctagagcgtatgctttggccaaacgtcctgggttgggagtggggtatacagactgacttttcttgaagtcttcgggatg
ctgggggagggggagggtcggacatcatatacatctccacccacagtgatggggaccaaacttccaggctagttgtggtttatgactTTgaagatggccgctcct
gagtatccgtgcctggtctctggtatttctgtgatgggatcctacagggacagcccctgacactgagtactgtgttgcccccagtatacagaggagaaaactgagagac

gggtaattgacgacagaccattcctggactggagaggtgggccttttaactgtccatcctgcatcaatttgaaatggatgacagagaggaaacttctttgcttctctgacc
acaactacttccaggaagagggtggggcagagaggccagggctccccgagtttttTTaaggagtggtggttctgcaagccatcaagcatttcagagggatgatggg
ccatgctggcttgacttttccctcctctccccactctgccttgtgtactttttggagccagaaagctgctttgtggtctgtggtggcaggggcctgggtgctgctggtggcttgg
gcaattccctttgcctctgggcctctgaagcaatggtgctgttaggagcccagctggaagggctgttccaagacacaagcagggcactagaaaggacccagcactg
gctctgagacagacgcctggagcttgctggtcatcaggggcccacatcccacaccatcttcatgagttctgtgactcctgcagcctgggacatgcttctagtggctttccta
atgcttaatgatcacgttcagaggaagtggctgctctgggcacactgacactgtctgggctttctctggagcgttagaatggctgtggttctccatcccagcatggccagtc
cttcagggaggtgaggagagagcatggaggaccaaacctgctgtgtgaatgaataaggttctccccatgctgcttttcagtctagcagggcgaagggcagagacctg
gtgagtactgtggtaccatgcagatccccacagcaagagcttgggggtggagggaggaggcagctagctgggaaacccctgcttgacacctggacccgccaggg
agttaagcccctgcctttggggaacaggctggccagaaggcaaagaaaagctcttggggtgaggcctcaccgttgcctggcttgtactggtactcaccagttccaggt
cacctttaaaacagttgcccttcacctgggcttctggttggccctgtggcactgctggcatccctgtttcacaggctggaaccaggccctccttcagacatggtgaataga
gaagctgttcttcagcatgccagggggatttcctatctgaagttgacaggtcctgttactcacacatctaaatcacagatccgctgttctgtaaaagcctgaggtagaaca
gctaggaataaaggctcccgtcacactccctggagctggccccttcttccctttgtgagtcattaaaaaaaaaaaaaaaaaaaaatccatcctgacatatggaaggtg
acctaaaagcccctgggagctacacttcagtgtctgtgtgtctgtggccctgtggcttcccgggggtacagattctctatctggcctcaaacctcagcctcatgattcagca
gagttgactgacgtcagcgcacaagaatgaaaagtgagaggagggctcaagtagggggccaggtggggtgtggagctggccgactgggaaaggaacagggct
gcttgcacacgcttagaagccgccgcagctggcaactgtggtccttgttttactgaggaggaagaggctcaggccttgaaggaggcctgacctgttttcagcaagatga
atttattatttttttcttttggtttttgagacagggttcctctgtgtatccccagctatcctggaactcactctgtagaccaggctgcctctgcctggtgcacctctggagtgcacta
ccactcctgtgaaagtacattgctaggctcccctcaccccgtaaactctcagcctagggagtttagttaggtgtgggatgggcaggcgagcctcttctatggggtgtccag
cacaaagcacttagcacagagacttgatacagtagaggcaacaggtacacaatgaaggacgtgcagagagttcaaggtctgcctgggctacatagtgagaaata
agaaacactgcctgtttccgatgttcttctaagccccaagaacacagaaataaatggaaataaatattctcttggaggataacagacaaatataaaggaacactcctct
ttaagaagaggcagtgctcagaggaagactaatggtggttggatgtgggcagcactgga

**Supplementary Table 3. Primers used in this study.**

| Mutagenesis primers (Seq 5'-3') | | |
|---|---|---|
| Human SOCS1 R104A Quickchange | CCTTCCTGGTGGCCGACAGCCGCCAGCGG | |
| | CCGCTGGCGGCTGTCGGCCACCAGGAAGG | |
| Human SOCS1 F58A Quickchange | GCACTTCCGCACAGCCCGTTCGCACGCCG | |
| | CGGCGTGCGAACGGGCTGTGCGGAAGTGC | |
| **Genotyping primers (Seq 5'-3')** | | PCR size |
| Mouse Socs1 NGS* | GTGACCATGAACTCAGGAGTCCGACACTCACTTCCGCAC | 297 bp |
| | CTGAGACTTGCACATCGCAGCAGCAGCTCGAAAAGGCAGTC | |
| Mouse Socs1 WT allele | CCTCCAGCTGGCCCCTCGAGTAGGATG | 394 bp |
| | CATTCGCCATTCAGGCTGCGCAAC | |
| Mouse Socs1 null allele | GCATCCCTCTTAACCCGGTAC | 185 bp |
| | AAATGAAGCCAGAGACCCTCC | |
| Mouse Ifng WT allele | CCTCAGAACTCAAGTGGCATAGAT | 266 bp |
| | TTCAATGACGCTTATGTTGTTGCTG | |
| Mouse Ifng null allele | CATTCGACCACCAAGCGAAACATC | 1000 bp |
| | TTCAATGACGCTTATGTTGTTGCTG | |

*NGS= next generation sequencing
